# Effect of error and missing data on population structure inference using microsatellite data

**DOI:** 10.1101/080630

**Authors:** Patrick A. Reeves, Cheryl L. Bowker, Christa E. Fettig, Luke R. Tembrock, Christopher M. Richards

## Abstract

Missing data and genotyping errors are common in microsatellite data sets. We used simulated data to quantify the effect of these data aberrations on the accuracy of population structure inference. Data sets with complex, randomly-generated, population histories were simulated under the coalescent. Models describing the characteristic patterns of missing data and genotyping error in real microsatellite data sets were used to modify the simulated data sets. Accuracy of ordination, tree-based, and model-based methods of inference was evaluated before and after data set modifications. The ability to recover correct population clusters decreased as missing data increased. The rate of decrease was similar among analytical procedures, thus no single analytical approach was preferable. For every 1% of a data matrix that contained missing genotypes, 2–4% fewer correct clusters were found. For every 1% of a matrix that contained erroneous genotypes, 1–2% fewer correct clusters were found using ordination and tree-based methods. Model-based procedures that minimize the deviation from Hardy-Weinberg equilibrium in order to assign individuals to clusters performed better as genotyping error increased. We attribute this surprising result to the inbreeding-like nature of microsatellite genotyping error, wherein heterozygous genotypes are mischaracterized as homozygous. We show that genotyping error elevates estimates of the level of genetic admixture. Overall, missing data negatively impact population structure inference more than typical genotyping errors.

## INTRODUCTION

Short, repetitive regions of the genome, known as microsatellite DNA, simple sequence repeats (SSRs), or short tandem repeats (STRs), are commonly used in molecular population genetic studies (Sunnucks 2000; Guichoux *et al.* 2011). SSR loci exhibit a unique mutational mechanism, slipped-strand mispairing, which causes the duplication or deletion of repeat units, resulting in sequence length variation among alleles (Levinson & Gutman 1987). SSR mutation rates vary widely depending on organism, repeat length, and repeat number, but are generally 1E3–1E4 times higher than a typical nucleotide substitution rate of 1E-8 per generation (Dallas 1992; Chakraborty *et al.* 1997; Vigouroux *et al.* 2002), thus SSR regions provide highly polymorphic markers, useful for distinguishing individuals, reconstructing population history, and estimating demographic parameters.

While single nucleotide polymorphisms (SNPs) may gradually supplant SSRs for certain population genetic applications (Brumfield *et al.* 2003), SSRs remain popular. Over 3500 papers that utilized SSRs were published in 2009 (Guichoux *et al.* 2011), and in 2013–2015, ~33% of articles in the journal *Molecular Ecology* included SSRs as a primary data source. SSR development and typing costs have dropped due to next generation sequencing (Gardner *et al.* 2011) and improved multiplexing protocols (Butler 2005; Holleley and Geerts 2009). We expect SSRs to remain in use due to low cost and their ability to outperform SNPs with fewer loci for individual identification (Seddon *et al.* 2005); parentage and sibship analysis (Glaubitz *et al.* 2003; Wang & Santure 2009); and population structure inference (Liu *et al.* 2005; Glover *et al.* 2010).

Most SSR data sets contain missing data and erroneous genotypes. Missing data are entered into an SSR data matrix when a particular sample does not produce an interpretable pattern of DNA fragments after PCR amplification. PCR failure is usually caused by poor quality template DNA or improper PCR conditions, including mispriming due to mutations at primer binding sites (Guichoux *et al.* 2011). The frequency of missing data is elevated across loci for samples with poor quality DNA. Suboptimal PCR conditions increase missing data across samples for specific loci. Consequently, missing genotypes typically occur non-randomly in data matrices, clumped in rows or columns. Missing data can be minimized by re-extracting DNA and repeating amplifications for problematic individuals, redesigning troublesome primers and optimizing PCR conditions, or excluding individuals and loci with high failure rates from the matrix.

SSR genotyping errors arise from three main sources: the occurrence of null alleles at a locus, preferential amplification of small DNA targets during PCR, and stuttered visualization of amplification products (DeWoody *et al.* 2006; Guichoux *et al.* 2011). “Null alleles” are genotypic variants that fail to amplify under the conditions specified for the locus. They often occur due to mutations at primer binding sites that inhibit amplification but may also arise from poor template quality. Because nothing is amplified, null alleles, by definition, cannot be observed and are not scored. Consequently, in diploid organisms, the genotype entered into the data matrix becomes—erroneously—homozygous for the other, visible allele, or, if both alleles are nulls, missing data. Several analytical approaches have been devised to detect null alleles and estimate their frequency (Chakraborty *et al.* 1992; Raymond & Rousset 1995; Van Oosterhout *et al.* 2004; Kalinowski *et al.* 2007) although only about 40% of studies use them (Guichoux *et al.* 2011). In most cases, eliminating null alleles requires primer redesign outside of highly mutable regions (Dakin & Avise 2004; Chapuis & Estoup 2007).

Preferential amplification of short DNA sequences causes “large allele dropout” error. All else equal, short sequences are more efficiently amplified than long. In a heterozygote with differently-sized alleles this bias may prevent the signal for the large allele from rising above the detection threshold, with the consequence that a heterozygous genotype will be erroneously scored as homozygous for the smaller allele (Wattier *et al.* 1998; Björklund 2005). Large allele dropout can be mitigated in some cases by excluding loci where amplicons exceed 200 bp (Sefc *et al.* 2003).

Slipped-strand mispairing during PCR results in the production of shadow peaks around the amplified allele (Murray *et al.* 1993), a phenomenon termed “stutter”. Stutter peaks are usually smaller than the target, and deviate in size by multiples of the repeat unit length, with progressively decreasing signal (Shinde *et al.* 2003). As with other types of error, stutter causes heterozygotes to be scored as homozygous, but always for the larger of the two alleles, and only when the alleles differ in size by a single repeat unit. Stutter can be reduced by avoiding SSRs with dinucleotide repeats (Chambers & MacAvoy 2000), decreasing denaturation temperature (Olejniczak & Krzyzosiak 2006), and using highly processive polymerases (Davidson *et al.* 2003).

The final assembly of an SSR data set requires considerable care. Filling all cells in a data matrix is time consuming; poor-performing loci must be optimized and recalcitrant individuals must be extracted and genotyped repeatedly. Consequently, the typical course for dealing with missing data is to eliminate problematic individuals or loci. This can produce biased sampling because the frequency of missing data may be similar in related populations (Amos 2006). Some authors have recommended that error rates be reported and efforts made to assess the reliability of conclusions given uncertainty in the genotypes (Bonin *et al.* 2004; Broquet & Petit 2004; Hoffman & Amos 2005). This practice is increasing—error rate estimates can be found in about a quarter of published papers (Guichoux *et al.* 2011)—but it requires sample replication and extensive post-genotyping data analysis, increasing costs. If missing data and genotyping error were to impact the accuracy of population structure inference in only minor ways these steps might be avoided.

The effect of missing data and error on linkage mapping (Hackett & Broadfoot 2003), parentage analysis (Dakin & Avise 2004; Hoffman & Amos 2005; Kalinowski *et al.* 2007), and the estimation of population genetic parameters (Chapuis & Estoup 2007; Hall *et al.* 2012; Peel *et al.* 2013) has been studied in detail, but there have been few investigations of the effect on population structure inference (Pompanon *et al.* 2005; Carlsson 2008; Chapuis *et al.* 2008). In this study we seek to quantify the extent to which missing data or genotyping error impact population structure inference, to assist researchers in developing strategies to produce SSR data sets that maximize accuracy while minimizing costs.

## MATERIALS AND METHODS

Highly polymorphic, neutral marker data were simulated using a coalescent model and the software MSMS (Ewing & Hermisson 2010). The simulation approach is detailed in Reeves *et al.* (2012). Briefly, 1E4 data sets were generated, each containing 500 diploid individuals equally distributed among 50 populations, and 50 unlinked loci. An asymmetric island migration model was created by randomly assigning migration rates between populations. The population scaled mutation rate (θ) was varied from 0–0.5. The underlying mutation model of MSMS is an infinite sites model. Unique binary strings output by MSMS were converted into uniquely named alleles following Huelsenbeck and Andolfatto (2007), rendering the infinite sites model as an infinite allele model. We used an infinite allele model instead of a stepwise mutation model as an expedient, and because it is not clear the extent to which the stepwise mutation model fits real SSR data (Gaggiotti *et al.* 1999). It is not unusual for SSR data sets to better fit an infinite allele model than stepwise mutation (e.g. Estoup *et al.* 1995; O’Connell *et al.* 1997). The primary defining feature of real SSR data is a limited number of alleles per locus (Paetkau *et al.* 1997). Therefore, data sets containing levels of polymorphism comparable to typical SSR loci (on average < 30 alleles per locus, following Kalinowski 2002) were subsampled from the original 1E4 data sets. A total of 1367 were found and used for further analysis.

Simulated data sets were altered to include missing or erroneous genotypes using models describing the distribution of these data aberrations in typical SSR data sets. In the missing data model, the percent of the matrix containing missing data was varied between zero and 25. A “clumping” parameter was used to bias placement of missing genotypes towards certain loci or individuals. The clumping parameter was varied among data sets from zero, which caused a uniform distribution of missing data, to ten, which elevated the probability ten-fold that the next missing genotype would occur in a row or column already containing missing data. Missing genotypes were substituted for known genotypes one by one, with the probability of conversion for each row and column adjusted after each substitution using the clumping parameter, until the specified percentage of missing data was reached.

To create data sets that varied in their propensity to contain null alleles, a data-set-specific maximum null allele frequency parameter (ν_d_) was defined, and selected at random from 0 to 20 percent, following Dakin and Avise (2004). Within each data set, the locus-specific null allele frequency parameter (ν_l_) was chosen at random from 0 to ν_d_ for each locus. The number of null alleles per locus was then defined as ν_l_ multiplied by the number of distinct alleles at the locus, with the alleles that were to act as nulls chosen randomly. Alleles defined as nulls were treated as unknown, with the consequence that heterozygous genotypes became homozygous for the non-null allele, and homozygous null genotypes became missing data, in the modified data set.

To simulate large allele dropout, it was necessary to assign a probability of dropout for each allele that was proportional to its size. A locus-specific maximum probability of dropout (δ_l_) was chosen at random for each locus from 0 to δ_d_, the data-set-specific dropout probability. The ceiling on δ_d_ was set to 0.5 based on empirical studies (Taberlet *et al.* 1996; Gagneux *et al.* 1997; Buchan *et al.* 2005). We assumed a curvilinear function relating dropout probability to allele size, with the largest allele at each locus having a dropout probability of δ_l_. Coalescent simulations only provide information on allelic state, not allele size, so relative allele sizes (σ_a_) were assigned to all alleles for each locus by randomly sampling an exponential distribution with rate parameter λ, varied by locus from 0 to 10. The probability of retention (i.e. the probability that an allele does not drop out) was then computed for each allele as one minus the cumulative distribution function of the exponential (CDF = 1 − e−^λσ_a_^) rescaled to have a maximum value of δ_l_. In this way, data sets exhibited varying levels of overall “dropout proneness” governed by the parameter δ_d_. Within data sets, the largest alleles at a locus always had the highest dropout probability, but some loci were characterized by dropout probabilities that declined uniformly with consecutively smaller allele size (when λ→0), while others approximated a threshold effect where alleles above a particular size were highly dropout prone (when λ→10). If a uniform random number from 0 to 1 exceeded the probability of retention, the allele was made to drop out, and the data set was modified accordingly. Thus, the effect of simulated large allele dropout was to convert heterozygous genotypes to small allele homozygotes with probability δ_l_(1 − e−^λσ_a_^).

To model stutter error, we identified all heterozygotes having consecutively-sized alleles using the arbitrary sizes from the large allele dropout model. These genotypes are called “adjacent-allele heterozygotes” (Hoffman & Amos 2005). Stutter error could only affect these genotypes, but was not assumed to affect all of them. A model was developed to assign a “probability of stutter error occurrence” to each adjacent-allele heterozygous genotype, thus permitting us to vary the “stutter error proneness” between data sets and among loci. A data-set-specific average probability of stutter error at adjacent-allele heterozygotes 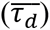 was randomly chosen from zero to one. 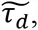 the standard deviation of 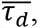 was randomly set from zero to one. The locus-specific probability of stutter error, τ_*l*_, was sampled from the normal distribution defined by 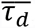 and 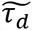. The conversion of adjacent-allele heterozygotes into large-allele homozygotes then occurred with probability τ_*l*_, the locus-specific conversion probability.

Genotypes in the unmodified matrices were altered in the order that errors would arise during the genotyping process. Null allele errors, which are caused by mispriming, were added first, followed by large allele dropout errors, caused by poor amplification, then by stutter errors, caused by slippage during amplification but attributable primarily to poor scoring procedures. Realized error rates were then recalculated for each error type, for each data set.

Three categorically-distinct analytical approaches for inferring population structure were used. First, we applied a class of Bayesian Markov chain Monte Carlo (MCMC) methods introduced by Pritchard *et al.* (2000) in the software STRUCTURE. To avoid *ad hoc* model selection procedures for determining the number of populations (K) (Evanno *et al.* 2005), we used INSTRUCT v1.0 (Gao *et al.* 2007) and STRUCTURAMA v1.0 (Huelsenbeck and Andolfatto 2007) instead of STRUCTURE. For INSTRUCT analyses, we used no-admixture (mode 0) and admixture (mode 1) models, as well as a model that estimates individual inbreeding coefficients simultaneously with individual assignment (mode 5). Modes 0 and 1 of INSTRUCT are comparable to no-admixture and admixture models of STRUCTURE. A single Markov chain was run for 1E5 generations and sampled every 25, with the initial 12500 generations discarded. The deviance information criterion (DIC) was used to determine the best value of K between 1 and 50 (Gao *et al.* 2011). A discrete assignment was created by assigning individuals to clusters based on the highest assignment probability in the Q-matrix. STRUCTURAMA estimates K alongside individual assignment. Chains were run as for INSTRUCT, the prior on number of populations was set to two, and no admixture was allowed.

Second, we used a tree-based approach. Neighbor-joining (NJ) trees were constructed using NTSYS (Rohlf 2008). Inter-individual distances were computed using Lynch’s (1990) band sharing coefficient. Populations were counted as correctly inferred when they existed as a monophyletic group in the resulting tree. Third, an ordination method was applied. We used PCOMC, a procedure that couples ordination with cluster analysis to simultaneously determine population number and membership (Reeves & Richards 2009). Principal coordinate analysis was performed using NTSYS. Distance matrices, computed as for NJ, were double-centered prior to the calculation. Principal coordinate values were weighted according to their contribution to the total variance, then subject to the density clustering algorithm PROC MODECLUS (SAS Institute, Cary, NC).

Correct and incorrect clusters resulting from application of each analytical method to each data set were counted. Correct clusters were those that contained all 10 individuals that belonged to a single population as specified in the coalescent simulation model, and no others. Incorrect clusters contained some, but not all, members of a population in the model, or individuals from more than one population. The “performance ratio” was defined as the number of correct clusters resulting from analysis of a modified data set divided by the number of correct clusters in the unmodified data set from which it was derived. A performance ratio < 1 indicates that data modification reduces accuracy, while a performance ratio > 1 indicates improved accuracy. We used the partition distance of Gusfield (2002) as a second, less strict measure of accuracy for methods that produce partitions (INSTRUCT and STRUCTURAMA). The partition distance (*PD*) has been used previously for this purpose, and is useful because it can quantify partially correct matches between clusters, unlike the performance ratio (Huelsenbeck and Andolfatto 2007; Choi and Hey 2011). It is defined as the number of elements that must be moved between clusters to make one partition identical to another. We normalized the *PD* to the range 0–1 by dividing by the maximum possible *PD* (Charon *et al.* 2006), calling the result *PD*_*n*_. We define the “partition distance ratio” as 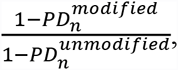, where numerator and denominator are formulated as similarities to simplify comparison with the performance ratio. 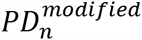 is the normalized partition distance between the partition resulting from analysis of the modified data and the 50 cluster partition defined in the coalescent simulation model (likewise for 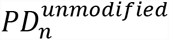). The performance ratio and the partition distance ratio are non-identical, but correlated, measures of the accuracy of population structure inference (Supplementary Table 1). We preferentially report the performance ratio, because its interpretation is intuitive, and it is applicable to all methods.

The false discovery rate (*FDR* was calculated as the number of incorrect clusters divided by the total number of clusters returned. A ratio, 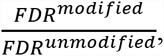 was calculated to express the difference in probability of recovering incorrect clusters between modified and unmodified data sets. Multiple regression and likelihood analysis were used to examine the effect of model factors on the accuracy of inference.

## RESULTS

### Data set validation

The 1367 simulated data sets contained an average of 14.48 ± 8.46 (1 sd) alleles per locus. Population structures ranged from virtually panmictic to highly subdivided (Figure 1). Levels of population subdivision, as quantified by Hedrick’s (2005) G’_st_, varied from 0.006 to 0.999, with an excess of low G’_st_ values. The degree to which data sets were modified by the missing data model was not significantly related to the level of population differentiation (*r* = 0.03, *p* = 0.32); application of the error models resulted in a slight, but significant, negative correlation between G’_st_ and the proportion of genotypes modified (*r* = −0.14, *p* < 0.0001 (Supplementary Figure 1).

**Figure 1.**
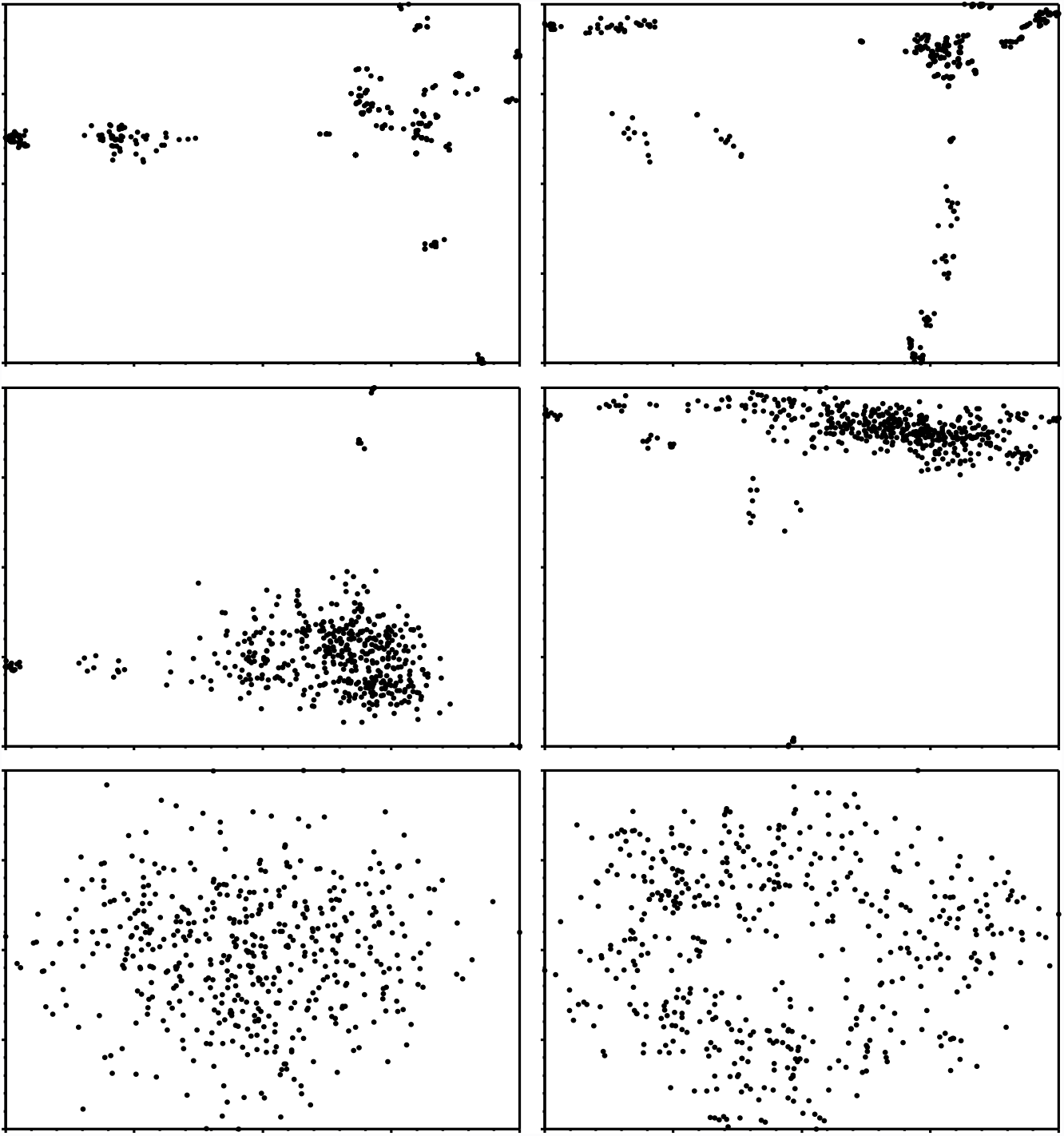
Some complex population structures generated by coalescent simulation, visualized using principal coordinate analysis. Top row, high population subdivision (G’_st_ = 0.99); middle row, intermediate (G’_st_ = 0.5); bottom row, low (G’_st_ = 0.01).

### Performance

The analytical methods differed in accuracy when unmodified data sets were analyzed (Figure 2). For a given level of genetic subdivision, correct clusters were found in NJ trees at a much higher frequency than in INSTRUCT, STRUCTURAMA or PCOMC analyses. This is not surprising because the criterion for identifying correct clusters for NJ was less strict than for other methods—a correct inference occurred when a node existed that defined a population correctly in the tree, but there was no mechanism to determine which nodes defined populations. PCOMC had the lowest rate of correct cluster recovery. STRUCTURAMA recovered more correct clusters and had a lower false discovery rate than INSTRUCT for high levels of genetic subdivision, while INSTRUCT was more accurate at lower levels, regardless of mode.

**Figure. 2.**
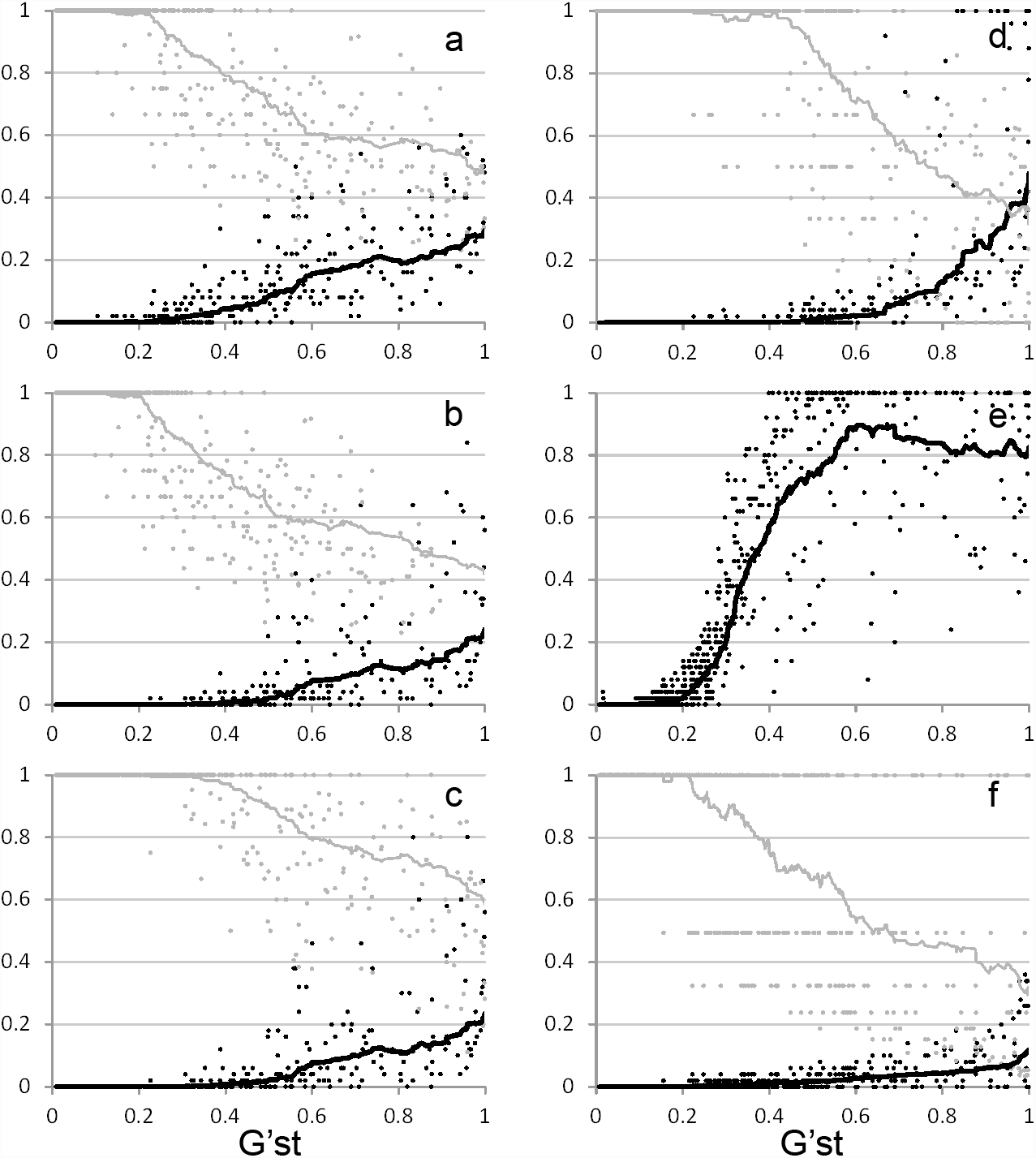
Accuracy of population structure inference for unmodified data sets. X-axis measures population subdivision, Y-axis shows the proportion of correctly inferred populations out of 50 possible (black), and the false discovery rate (grey). a) INSTRUCT no admixture; b) INSTRUCT admixture; c) INSTRUCT inbreeding; d) STRUCTURAMA; e) Neighbor-joining; F) PCOMC. False discovery rate not calculable for neighbor-joining.

To avoid undefined values when calculating the performance ratio, data sets were excluded when zero correct clusters were inferred with unmodified data. Likewise, for the partition distance ratio, data sets were excluded when 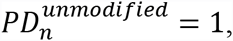, i.e. when the observed partition was maximally distant from the simulated partition. Because the coalescent model was complex and population subdivision was often low, a substantial number of data sets were excluded using this restriction. The total number of useful data sets ranged from 63 to 473, depending on analytical method, when measured using the performance ratio (Table 1), and 115 to 464 for the partition distance ratio (Supplementary Table 1). Population subdivision in the excluded data sets was low (G’_st_ ~5-fold lower than retained data sets), approaching or exceeding the limits of resolution of the methods applied.

**Table 1.**
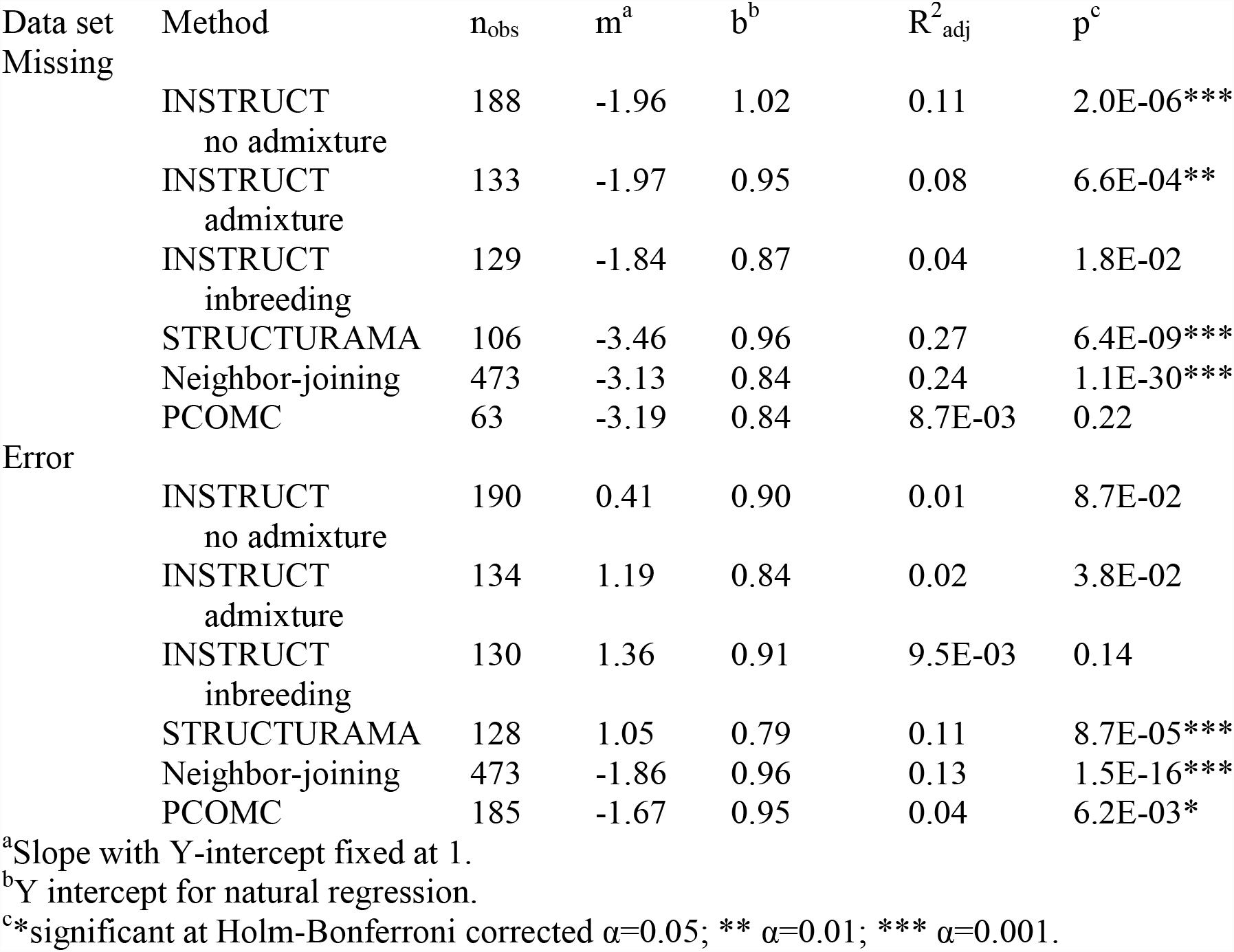
Regression analysis of clustering accuracy, measured with the performance ratio.

Performance decreased in a roughly linear manner as missing data increased (Figure 3a–f). The slope of the regression line was significantly different from zero (at α = 0.01, Holm-Bonferroni corrected, here and throughout) for all methods except ‘INSTRUCT inbreeding’ (mode 5) and PCOMC (Table 1). R^2^_adj_ values for significant regressions ranged from 0.08 to 0.27 and the slope of performance loss ranged from *m* = −1.8– −3.5 between analytical approaches (Table 1). The results using the partition distance ratio were similar. A significant negative correlation was found for ‘INSTRUCT no admixture’ (mode 0) and ‘INSTRUCT admixture’ (mode 1); the correlation was not significant for ‘INSTRUCT inbreeding’ or STRUCTURAMA (Supplementary Table 1).

**Figure. 3.**
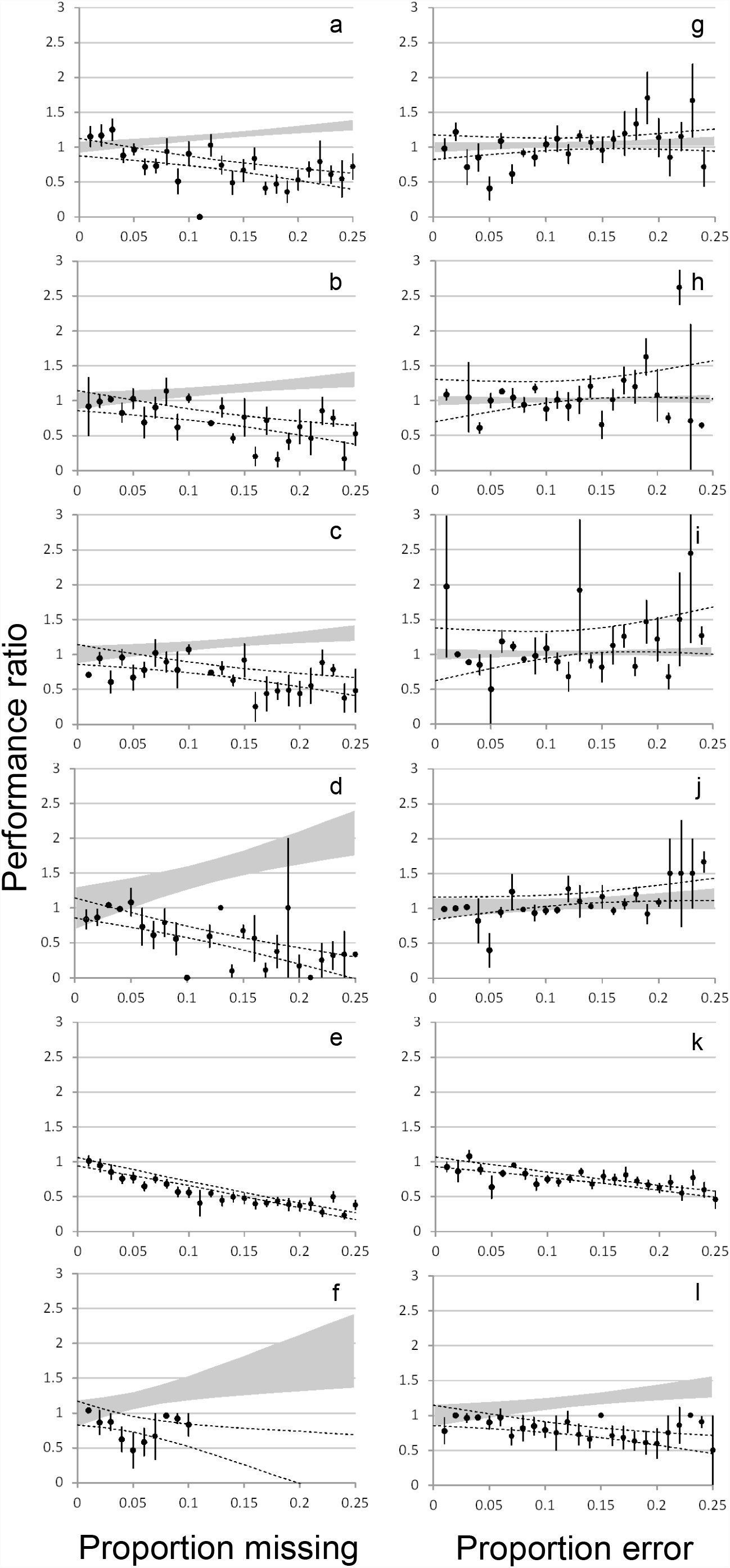
Effect of missing data and error on clustering accuracy using the performance ratio. a,g) INSTRUCT no admixture; b,h) INSTRUCT admixture; c,i) INSTRUCT inbreeding; d,j) STRUCTURAMA; e,k) Neighbor-joining; f,l) PCOMC. For visual clarity, points are mean values binned with an interval width of 0.01. Error bars indicate SEM. Statistics were calculated without binning. Dotted curves mark the 95% confidence interval on the slope of the regression. Grey shaded area is the 95% confidence interval on the slope of the regression for the false discovery rate ratio. False discovery rate not calculable for neighbor-joining. The range of values in f) is truncated because principal coordinate analysis can accept limited missing data.

Taking into account 95% confidence intervals on the slopes, a data matrix with 5% missing data is predicted to result in recovery of at least 72–81% of the correct clusters that could be recovered with a complete data matrix. Based on our data, 95% of recoverable clusters should be found when data matrices contain, for INSTRUCT, 2.5–2.7% missing data, or, for the other methods, 1.5–1.6% missing data. The FDR ratio increased significantly (α = 0.05) with missing data for all methods except ‘INSTRUCT inbreeding’ and PCOMC, indicating that, in addition to fewer correct clusters, users should expect more erroneous clusters as missing data increase.

The clumping parameter had a statistically significant effect on performance for NJ only (Table 2). Performance of NJ was higher when missing values were more clumped (*β* = 0.02). Using ratios of the Akaike weights, the linear model for percent missing data alone had a much higher probability (10^25^-fold) than the model for clumping parameter for NJ. Regardless of method, less than 4% of the variance in performance was attributable to clumping—a slight effect—thus the clumping parameter, despite adding realism to the simulation, was largely dispensable.

**Table 2.**
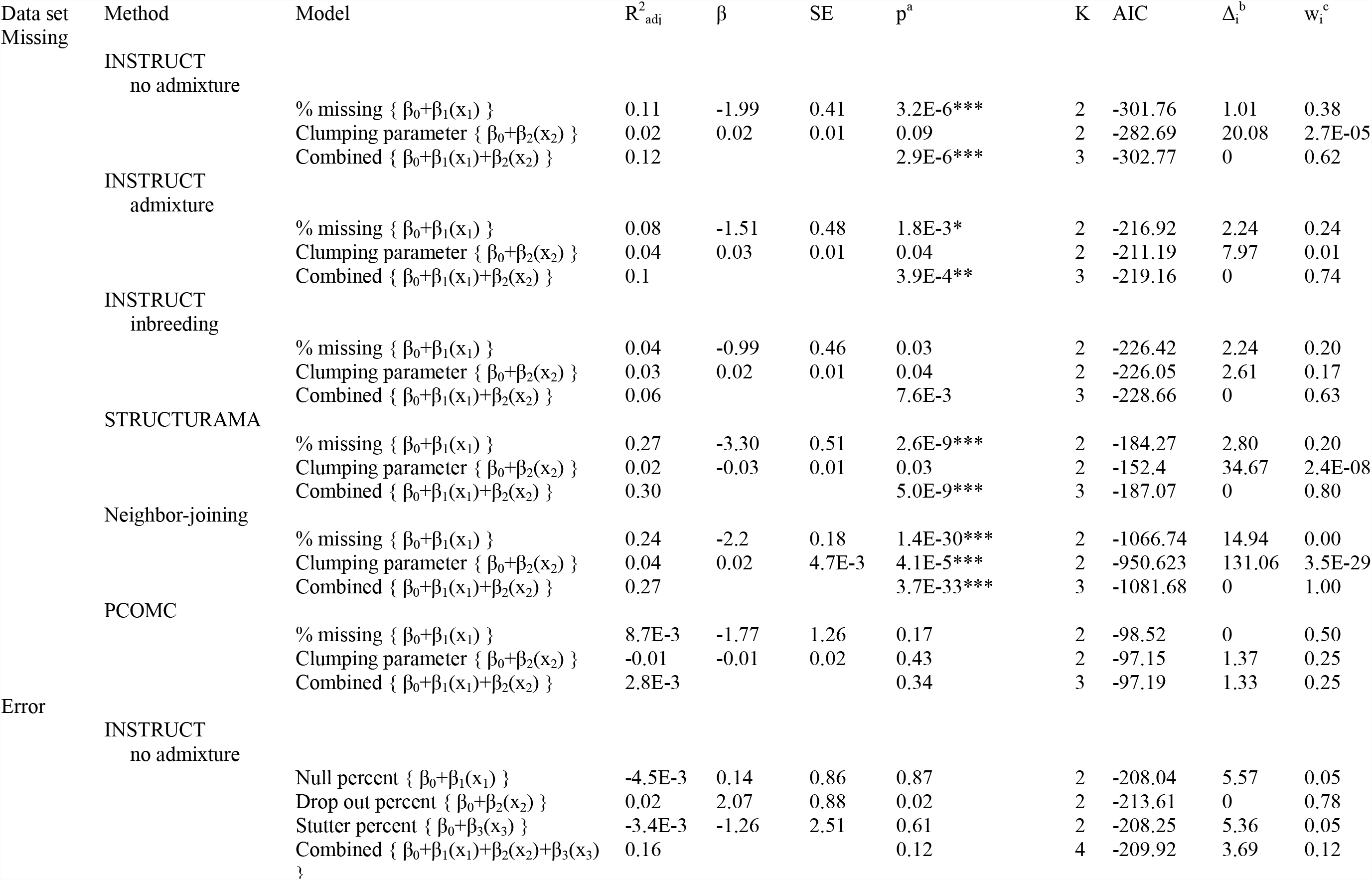
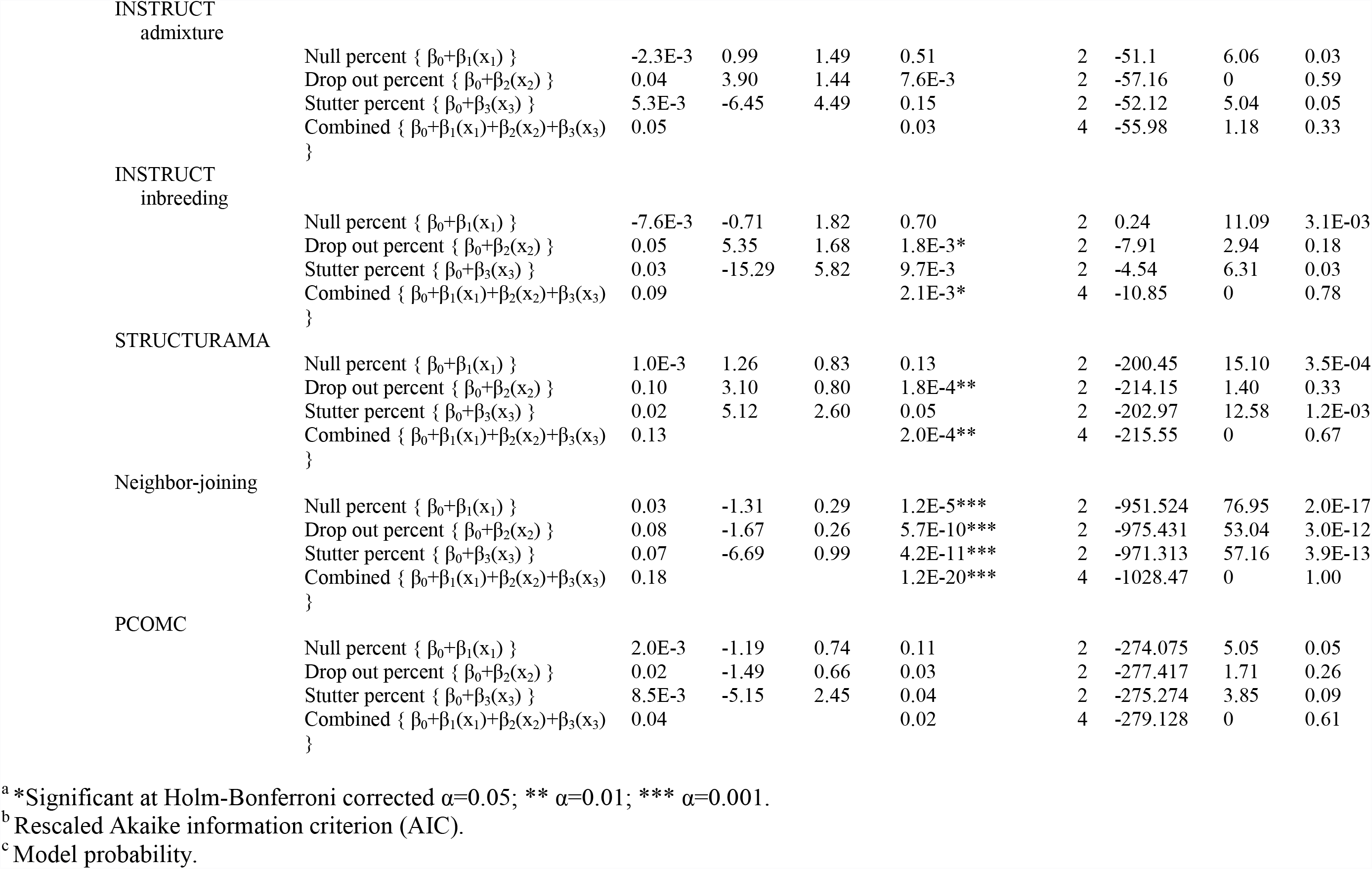
Multiple regression analysis and linear model likelihoods.

The effect of genotyping error differed between categories of analytical method. For distance methods NJ and PCOMC, performance declined as erroneous data increased (Figure 3k,l). The slopes of the regressions were significantly different from zero and about half the magnitude found for missing data (Table 1). A matrix with 5% erroneous data should result in recovery of 85% (NJ) or 82% (PCOMC) of the clusters that would be found if the data set contained no error. Ninety five percent of recoverable clusters were found when 2.9% (NJ) or 3.6% (PCOMC) of the data matrix was erroneous. For NJ, large allele dropout and stutter had the greatest effect on deteriorating performance (Table 2).

In contrast, accuracy of model based methods improved as erroneous data increased. When using the performance ratio, the slope of the regression was positive for all methods (0.41–1.36) (Figures 3g-j, Table 1). The relationship was statistically significant for STRUCTURAMA. When using the partition distance ratio, a significant positive correlation was found for all methods (Figure 4, Supplementary Table 1). Large allele dropout was the most important model effect for INSTRUCT and STRUCTURAMA, explaining 2–10% of the variation in performance (combined model, 5–16%), and holding 18–78% of the linear models’ Akaike weight (model probability). The FDR ratio did not change significantly with increasing error for any method.

**Figure. 4.**
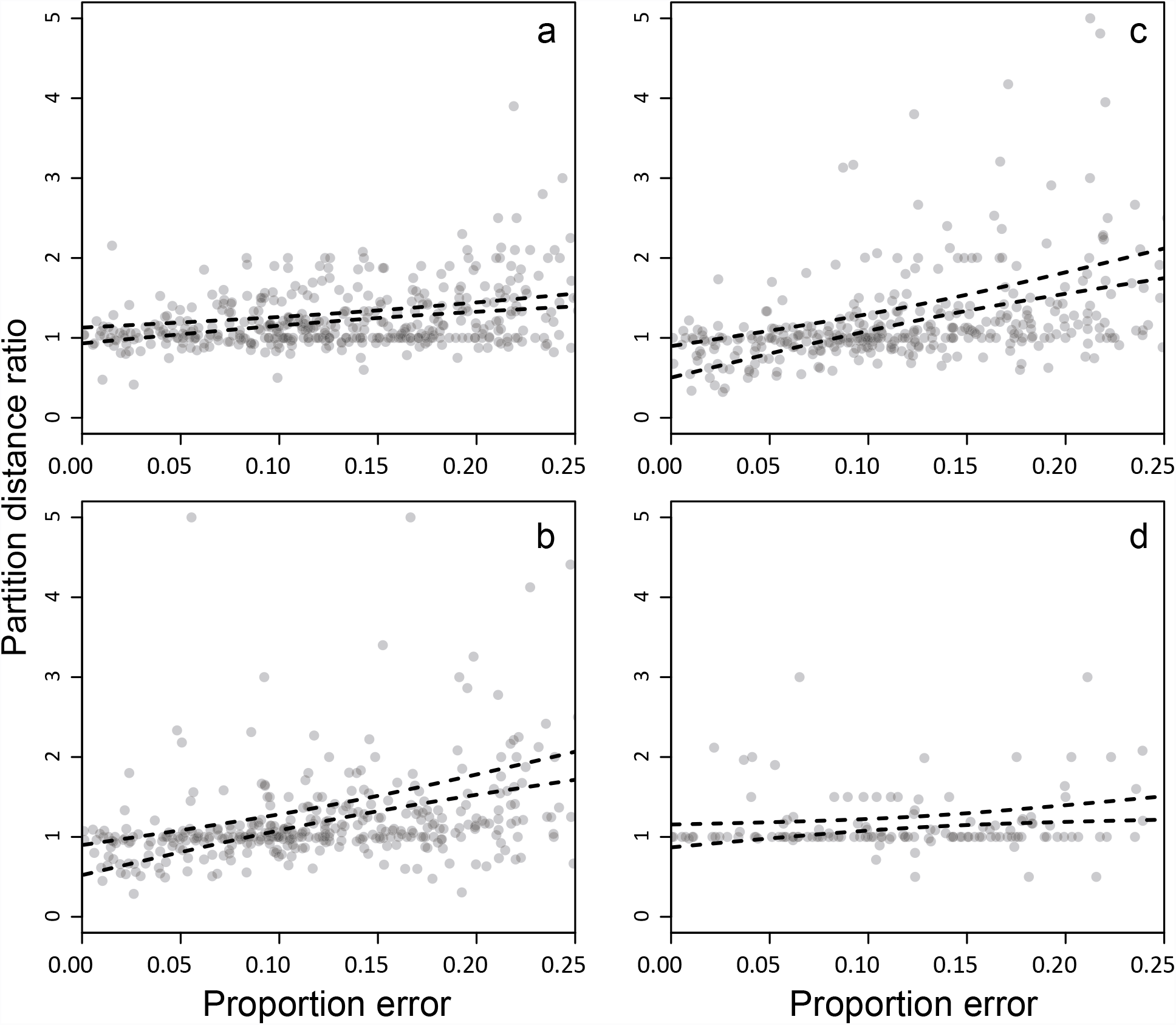
Effect of error on clustering accuracy using the partition distance ratio. a) INSTRUCT no admixture; b) INSTRUCT admixture; c) INSTRUCT inbreeding; d) STRUCTURAMA. Dotted curves mark the 95% confidence interval on the slope of the regression. For clarity, a total of 8 outlying points have been omitted from the plots, but were included in the regression.

## DISCUSSION

Genomewide genotyping approaches are gaining popularity for studies of population structure but SSRs continue to be used due to low cost and high power of inference. Although SSR data sets are prone to containing missing data and erroneous genotypes, few studies have examined their impact on analyses of population structure (Pompanon *et al.* 2005). We explored the effect of missing data and genotyping errors on population structure inference using model-based Bayesian MCMC procedures (INSTRUCT, STRUCTURAMA), a tree-based method (NJ), and an ordination approach (PCOMC). Our goal is to provide users of SSRs with insight for how much missing data and error might be tolerated in order to achieve a desired level of accuracy.

In order to make general recommendations it was necessary to explore a diverse set of population structures. This was accomplished using coalescent modeling with key parameters— mutation, migration rates, migration directionality—set stochastically, but within plausible ranges (Reeves *et al.* 2012). The resultant data sets exhibited a large range of complex population structures with widely varying levels of subdivision (Figure 1, Supplementary Figure 1). Nevertheless, the extent to which simulated data can ever accurately represent nature is debatable. We attempted to produce a realistic subsample from the universe of plausible population structures. Other models describing population divergence exist, most notably the isolation with migration model (Nielsen & Wakeley 2001). Our conclusions should be interpreted as limited to modified island models similar to those simulated, and should not be expected to precisely predict outcomes for any single data set.

Key properties peculiar to SSR data sets were modeled, including the clumped nature of missing data, and the three best understood sources of error: null alleles, large allele dropouts, and stutter artifacts. We did not include an assessment of human error, which contributes substantially (Bonin *et al.* 2004), but is not easily modeled. While most real data sets will be affected by both missing data and error, we elected to study these factors independently because the course of action for mitigation seems distinct.

### Impact of missing data

For data sets modified with missing data three basic observations were made. First, the degradation of performance as missing data increased was roughly linear (Figure 3a–f). Second, the rate at which performance declined was similar among methods (Table 1). Third, the degree to which missing data were clumped by locus or DNA sample was unimportant (Table 2)—contiguous blocks of missing genotypes did not affect performance much more than an equal number of randomly-distributed missing genotypes. These observations lead to a simple prediction: for every 1% of a data matrix that contains missing data, researchers should expect to recover 2-4% fewer correct clusters than would be inferred with a complete data set.

### Impact of error on distance methods

Error caused a decline in performance for the tree-based NJ method and for the ordination approach PCOMC. Researchers should expect a ~2% decrease in the number of correct clusters recovered for every 1% of the data matrix impacted by error when using these methods.

Although NJ and PCOMC are quite distinct mathematically, the similarity in response between the methods may arise from their use of the same distance metric. Distance was calculated as 1 − *S*_*xy*_, where *S*_*xy*_ is the ratio of alleles common to two individuals, over the total, or 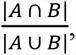 i.e. the Jaccard index (here, A and B represent the sets of unique alleles from each individual). The mechanism by which performance decreases is loss of precision in the genetic distance estimate. Conversion of heterozygotes to homozygotes decreases the number of alleles, lowering |*A* ∪ *B*|. The maximum number of distinct values the index can assume, |*A* ∪ *B*| + 1, also decreases, imposing a limit on the assembly of individuals into different clusters. Missing data had a stronger effect than error (Figure 3e,f,k,l). Conversion of a heterozygous locus to missing data may reduce |*A* ∪ *B*| by two, whereas with error |*A* ∪ *B*| can only decrease by one, thus the erosion of resolving power is generally more rapid with missing data.

### Impact of error on model-based methods

We observed a positive relationship between the amount of error introduced into a data set and the accuracy of inference when using model-based methods (Figures 3-4, Table 1, Supplementary Table 1). Our data suggest that a matrix containing 25% erroneous data could cause the recovery of 30–70% more correct clusters than an error-free matrix. Hereafter, we attempt to explain this unexpected result.

When a population consists of several subpopulations, a “heterozygote deficit” occurs, where the observed heterozygosity of the population analyzed as a whole is lower than predicted under Hardy-Weinberg equilibrium. This “Wahlund effect” (Wahlund 1928) forms the theoretical justification, and source of signal, for a class of model-based clustering methods that includes STRUCTURE, STRUCTURAMA, INSTRUCT, and others (e.g. Dawson and Belkhir 2001; Corander *et al.* 2003), which minimize the deviation from Hardy-Weinberg expectations by partitioning individuals into distinct subpopulations. Unfortunately, the signal upon which these methods rely is not uniquely generated by population subdivision. Heterozygote deficit also occurs as a consequence of selfing or inbreeding (Gao *et al.* 2007). Likewise, because SSR genotyping error typically involves the conversion of heterozygotes to homozygotes, it too induces a heterozygote deficit. In our simulation, a decline in observed heterozygosity with increasing error inflated the inbreeding coefficient 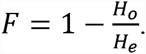. As error increased, the heterozygote deficit 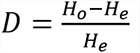 likewise increased (since D is the additive inverse of F). Thus error produces an inbreeding-like signature in data sets, which magnifies the Wahlund effect and, as a byproduct, inflates the level of genetic subdivision, causing increased signal (Supplementary Figure 2).

If this sort of signal amplification is important we would expect model factors ∆F (the change in inbreeding between unmodified and modified data due to error), and/or ∆G’st (the change in genetic subdivision) to drive the positive relationship between error and accuracy. Some evidence supports this. When accuracy was measured using the partition distance ratio, ∆G’st was the second-most important model effect for all INSTRUCT modes, explaining 3-10% of the variation (Supplementary Table 3). For ‘INSTRUCT no admixture’, ∆F was the most important. ∆G’st and ∆F were also important model effects for STRUCTURAMA, explaining 5-15% of the variation in the partition distance ratio. However, for STRUCTURAMA, the overwhelming weight of evidence was that ∆K, the difference in the number of populations inferred, was the primary driver of the response. Thus the improvement in accuracy observed for STRUCTURAMA seems to have an additional underlying cause.

STRUCTURAMA uses a Dirichlet process prior to simultaneously infer K and individual assignment. A “Dirichlet process” describes a distribution of probability distributions and is a mathematical construct commonly used as a prior in MCMC procedures to categorize items into groups when the number of groups is unknown. During progression of the Markov chain in STRUCTURAMA, a critical decision is made for each individual at each step: whether the individual belongs to an existing cluster or should be assigned to a new cluster. This decision affects both assignment and the inference of K. The algorithm proceeds roughly as follows. First, an individual *i* is selected at random from the existing partition and removed. Individual *i* is then re-assigned to whichever of the K clusters it fits best, or to a new cluster, all by itself. The probability that the individual is re-assigned to an existing cluster, *k*, is dependent on the number of individuals, η, in that cluster (large clusters are more attractive) and the marginal posterior probability of drawing *i*’s genotype from cluster *k*. In contrast, the probability of assigning *i* to a new cluster depends on the concentration parameter, α, of the Dirichlet process (the higher α is, the lower the probability that two randomly drawn individuals belong to the same cluster) and the probability of drawing *i*’s genotype from the prior distribution of allele frequencies, where all alleles are equiprobable.

In our unmodified data sets, the deviation from Hardy-Weinberg equilibrium is due to subpopulation structure plus some minor, random effects of the coalescent. However, when error is introduced, part of the deviation is then derived from the inbreeding-like effect of SSR error. With error, the relative probability of assigning an individual to an existing cluster versus a new cluster decreases—data with heterozygote deficit, by definition, do not fit nicely into a Hardy-Weinberg scenario—and, accordingly, it becomes more likely that an individual will be assigned to a new cluster, thereby increasing K. Consistent with this expectation, K increased significantly with error (Figure 5a). When more clusters are available in which to distribute individuals, more correct inferences can be made (equivalently, the number of correct inferences possible is suppressed for low levels of error). Consequently, performance improves with increasing error primarily because K increases. While convenient, the STRUCTURAMA approach for defining K may be incautious. Miller and Harrison (2014) argued that Dirichlet process models should not be used to infer the number of components.

**Figure. 5.**
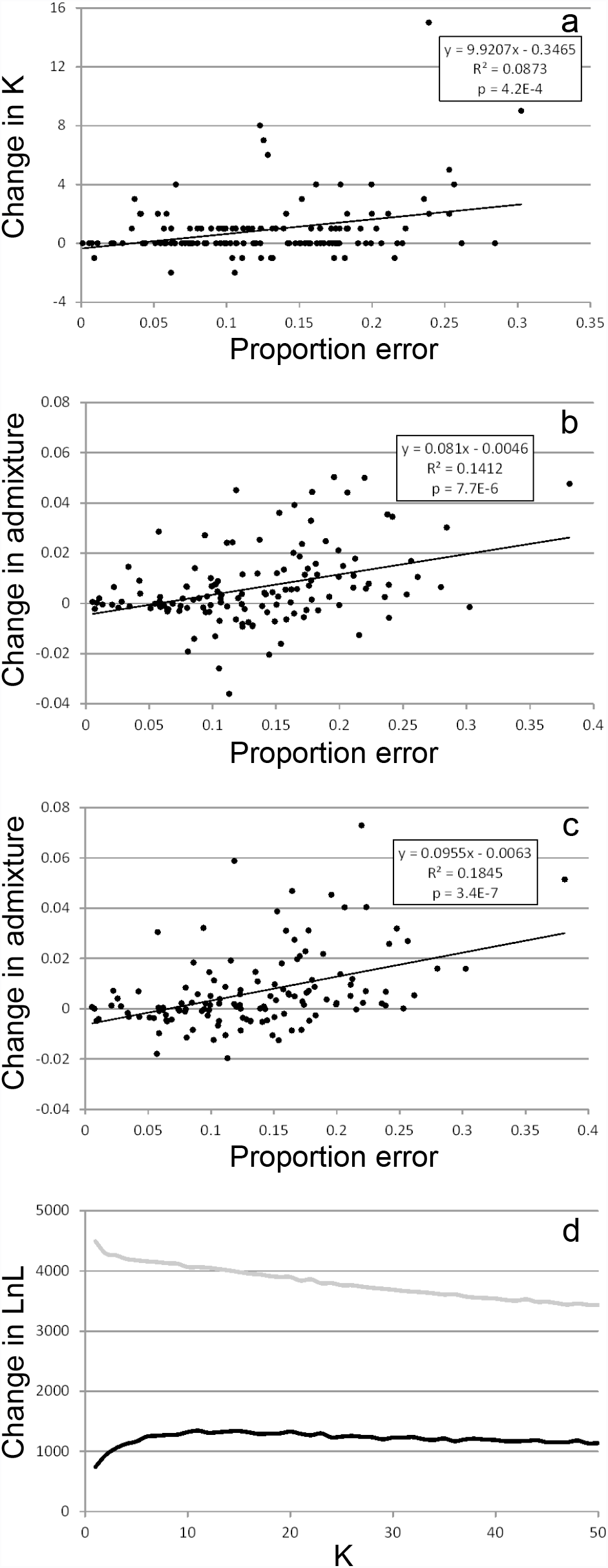
a) As genotyping error increases the optimal value for K is elevated under a Dirichlet process prior using STRUCTURAMA. b) The level of inferred admixture, as measured by the Shannon index, increases with error for ‘INSTRUCT admixture’. c) The upward bias in the admixture estimate is not corrected by using ‘INSTRUCT inbreeding’. d) Error causes the likelihood to be elevated for higher values of K, plateauing after K > 10. This effect is reversed by using a model that compensates for inbreeding. Black line, ‘INSTRUCT admixture’; grey line, ‘INSTRUCT inbreeding’.

INSTRUCT uses the Deviance Information Criterion (DIC) to select the best K value from a user-defined range. In the present case, a model with K=50 will usually have a higher likelihood than one with, say, K=4, but that does not necessarily mean it is better, it may merely be overfit. The DIC, like other model selection statistics, deals with this by imposing a penalty on the likelihood for adding parameters. The DIC value is, essentially, the mean likelihood across MCMC draws at stationarity, penalized for increasing K. The K value that minimizes the DIC across the range is deemed optimal. This method for choosing K appears largely insensitive to the level of error, and in this sense seems preferable to the Dirichlet process used by STRUCTURAMA. There is currently no consensus on the best procedure to estimate the number of groups in a finite mixture. Until the basic mathematical issues are resolved, the biological problem of estimating K for population structure inference will remain more craft than science.

### Impact of error on the estimation of admixture proportions

Using an admixture model and STRUCTURE, Gao *et al.* (2007) showed that inbreeding causes two spurious results: exaggerated levels of admixture and elevated likelihoods for higher K values. Falush *et al.* (2003, p. 1572) predicted this: “…situations that might cause additional populations to be inferred by STRUCTURE…include a significant frequency of inbreeding, cryptic relatedness within the sample, or the presence of null alleles.” The implication is that heterozygote-deficit-inducing factors other than population subdivision might promote the inference of more clusters, or more admixture among clusters, than is actually the case. For ‘INSTRUCT admixture’, the relative level of inferred admixture increased with error, and likelihoods were elevated for higher K values (Figure 5b,d). This did not, however, translate into the inference of higher K values under the DIC (*p* > 0.19).

To reduce artifacts from conflation of signal types, Gao *et al.* (2007) relaxed the assumption of Hardy-Weinberg equilibrium to estimate an inbreeding parameter alongside subpopulation membership. Relative to ‘INSTRUCT admixture,’ the relationship between error and admixture for ‘INSTRUCT inbreeding’ remained positive, but that between K and the model likelihood was reversed: *Ln*L decreased as K increased (Figure 5c,d). Thus the inbreeding model of Gao *et al.* (2007) compensates in some ways for genotyping error, but does not resolve the issue of admixture overestimation. The effects of error might be mitigated by explicitly modeling it as an additional parameter that alters Hardy-Weinberg expectations at each locus independently, distinct from the inbreeding coefficient estimated by INSTRUCT, which alters expectations uniformly across loci.

Whether over-estimation of admixture proportions due to genotyping error is a problem in real data sets is unknown, but the potential exists. False confidence in our understanding of the partitioning of genetic variation in nature notwithstanding, artifactually-elevated admixture estimates could have important economic implications, in endangered species management or genomewide association analysis, for example. Newer forms of data, like SNPs, are not immune to the problem. High throughput genotyping and next generation sequencing approaches vary in their capacity to accurately identify heterozygotes (Harismendy *et al.* 2009; Skotte *et al.* 2013), sometimes defaulting to the homozygous condition, akin to SSR error.

### Recommendations

Missing data exert a greater negative effect on the inference of population structure than typical SSR genotyping errors. As missing data increase, the number of correct clusters recovered decreases and the number of incorrect clusters increases. In order to efficiently produce SSR data sets for inferring population structure we recommend limiting the percent of a matrix that contains missing data to ~2%, unless a greater amount can be justified based on the particular hypotheses under examination. This will allow researchers to retain most of the resolving power of their data while not incurring the extra costs associated with completing a data matrix. For analyses reliant on simple distance metrics, we recommend limiting the error rate to ~4% of scored genotypes. Model-based population structure inference methods handle genotyping error well. We recommend the use of admixture models to infer cluster membership, but caution that admixture estimates may be artificially elevated.

## ACKNOWLEDGEMENTS

This study was conceived during a graduate seminar at Colorado State University organized by Mark P. Simmons, who we thank for critical reading of the manuscript. This research used resources from the Open Science Grid (supported by the National Science Foundation and the U.S. Department of Energy's Office of Science), the iPlant Collaborative (National Science Foundation grant #DBI-0735191. URL: www.iplantcollaborative.org), and the HTCondor group in the Department of Computer Sciences at the University of Wisconsin-Madison. We thank Nirav Merchant, Suchandra Thapa, Greg Thain, John Fonner and Nicole Hopkins for helping us find spare compute cycles and optimize HTC strategies at iPlant, the OSG, and UW.

## DATA ACCESSIBILITY

Computer programs used to simulate missing data and genotyping error are available from the authors.

## AUTHOR CONTRIBUTIONS

Designed research, all authors. Ran analyses, all authors. Analyzed results, PAR. Wrote computer programs, PAR. Wrote the manuscript, all authors.

## SUPPLEMENTARY FIGURE LEGENDS

Supplementary Figure 1. Properties of simulated data sets. a) Frequency of data sets with differing levels of population subdivision. b) The amount of missing data introduced into data sets was unrelated to level of population subdivision. c) The frequency of erroneous genotypes was negatively correlated with population subdivision.

Supplementary Figure 2. The inbreeding-like effect of SSR genotyping error. a) The level of apparent inbreeding increases with genotyping error. b, c) The increase in F is driven by a decrease in observed heterozygosity rather than an increase in expected heterozygosity, which decreased significantly, but slightly. d) Error increases the apparent level of population subdivision, producing a tendency for more accurate assignment of genotypes to clusters as the amount of error increases.

**Supplementary Table 1.**
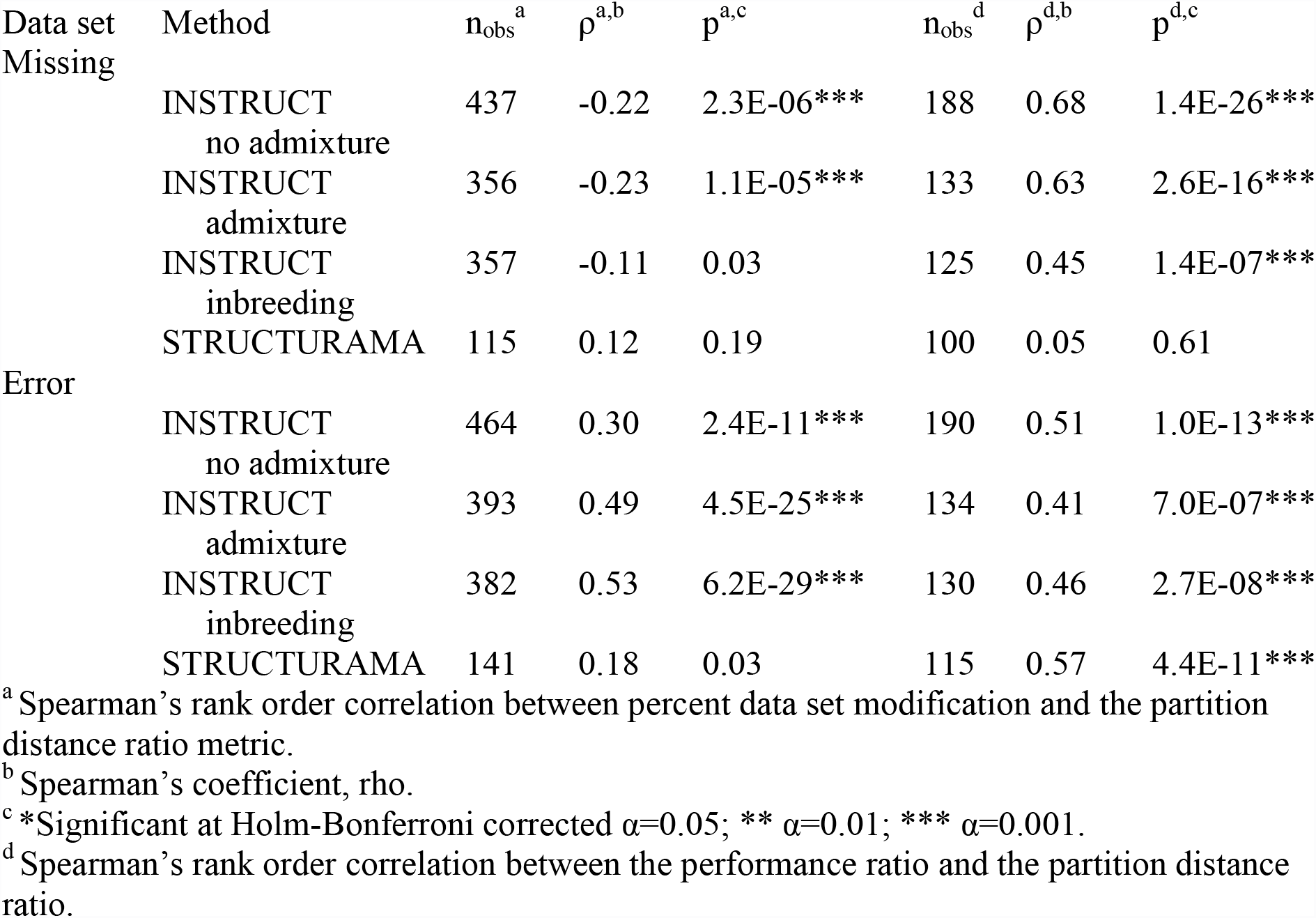
Spearman’s rank order correlation of clustering accuracy, measured with the partition distance ratio.

**Supplementary Table 2.**
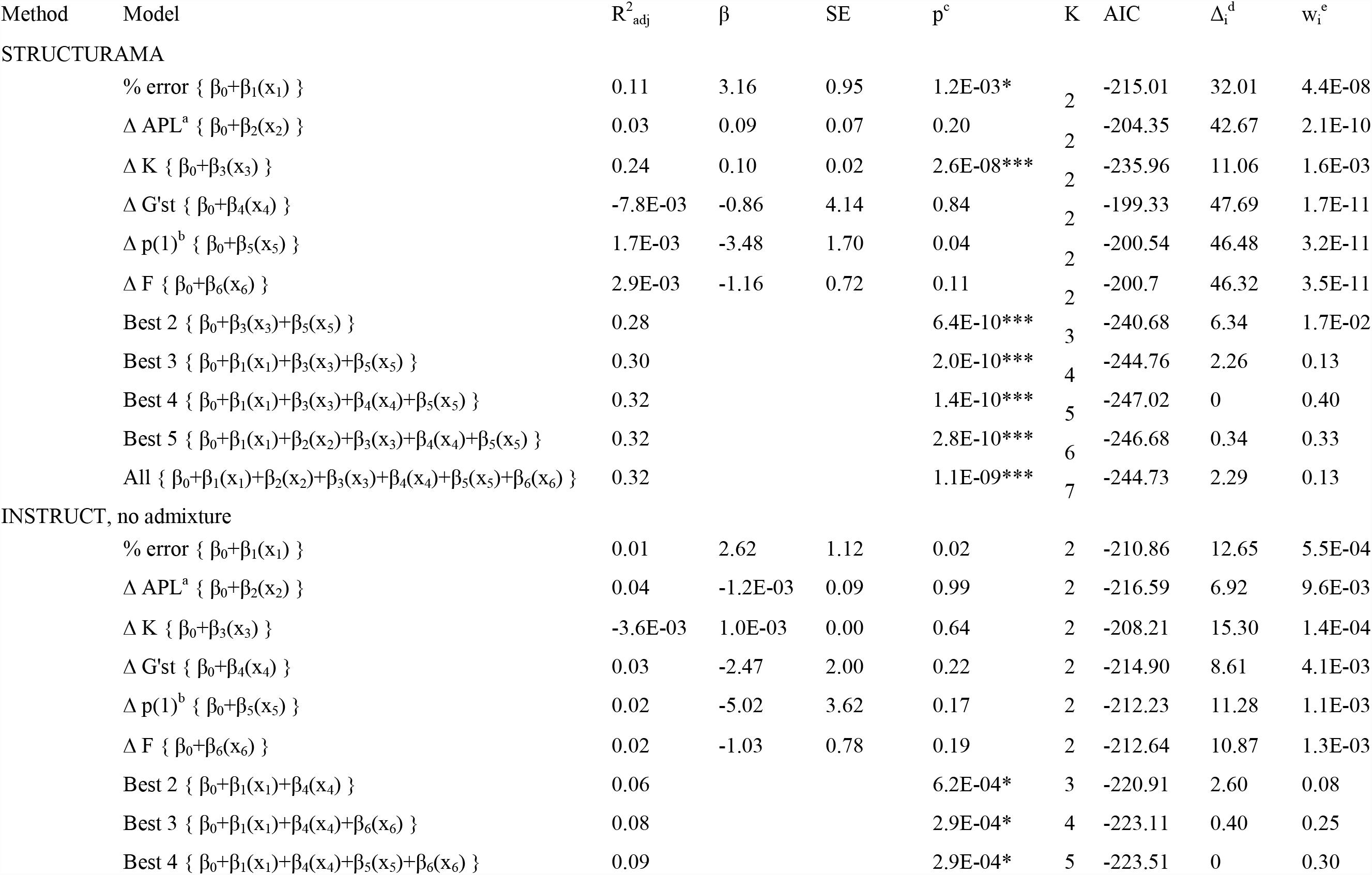
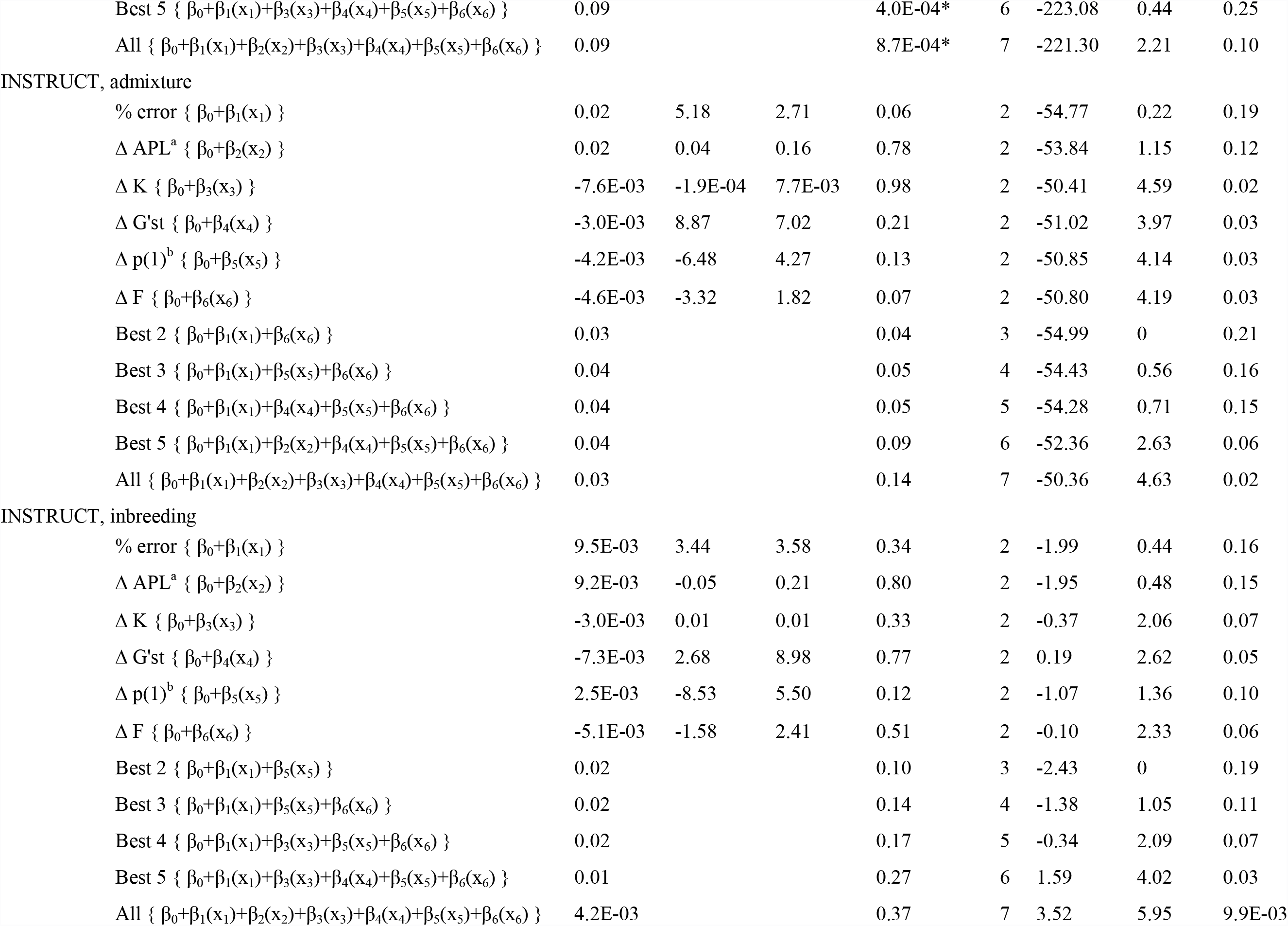
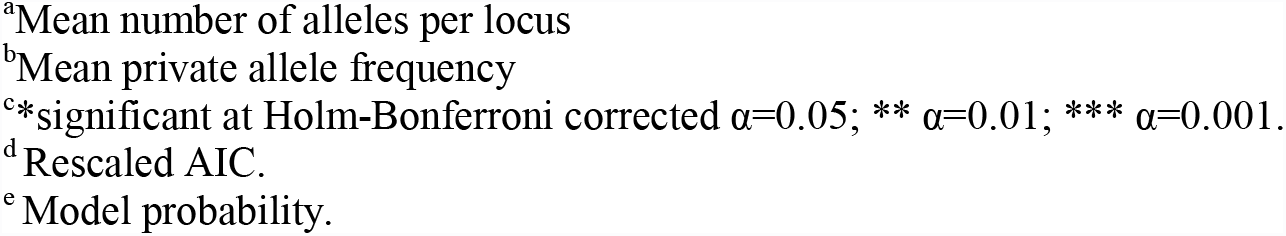
Multiple regression and likelihood analysis of six competing model effects explaining accuracy of model-based clustering methods subject to erroneous data, measured with the performance ratio.

**Supplementary Table 3.**
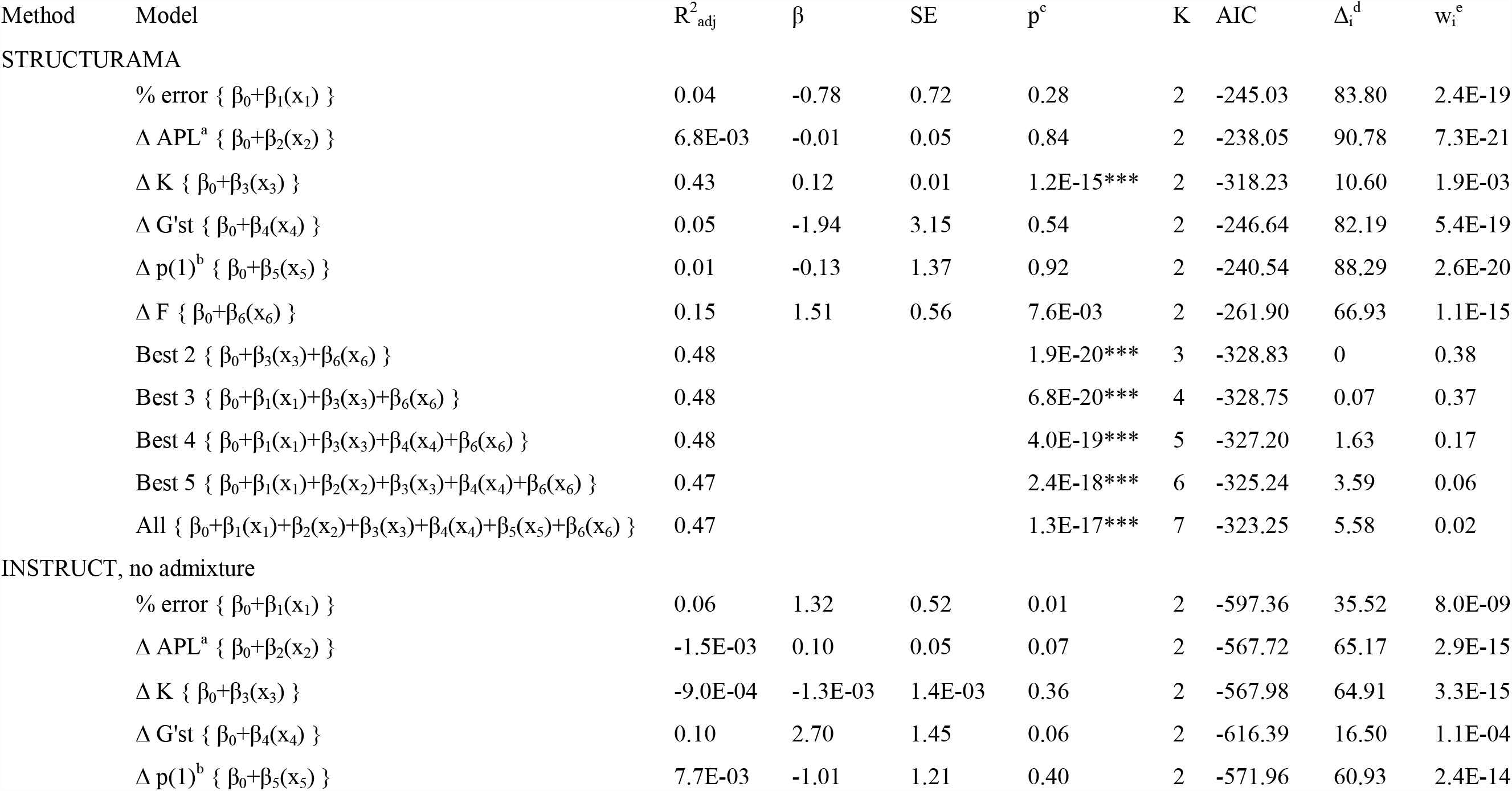
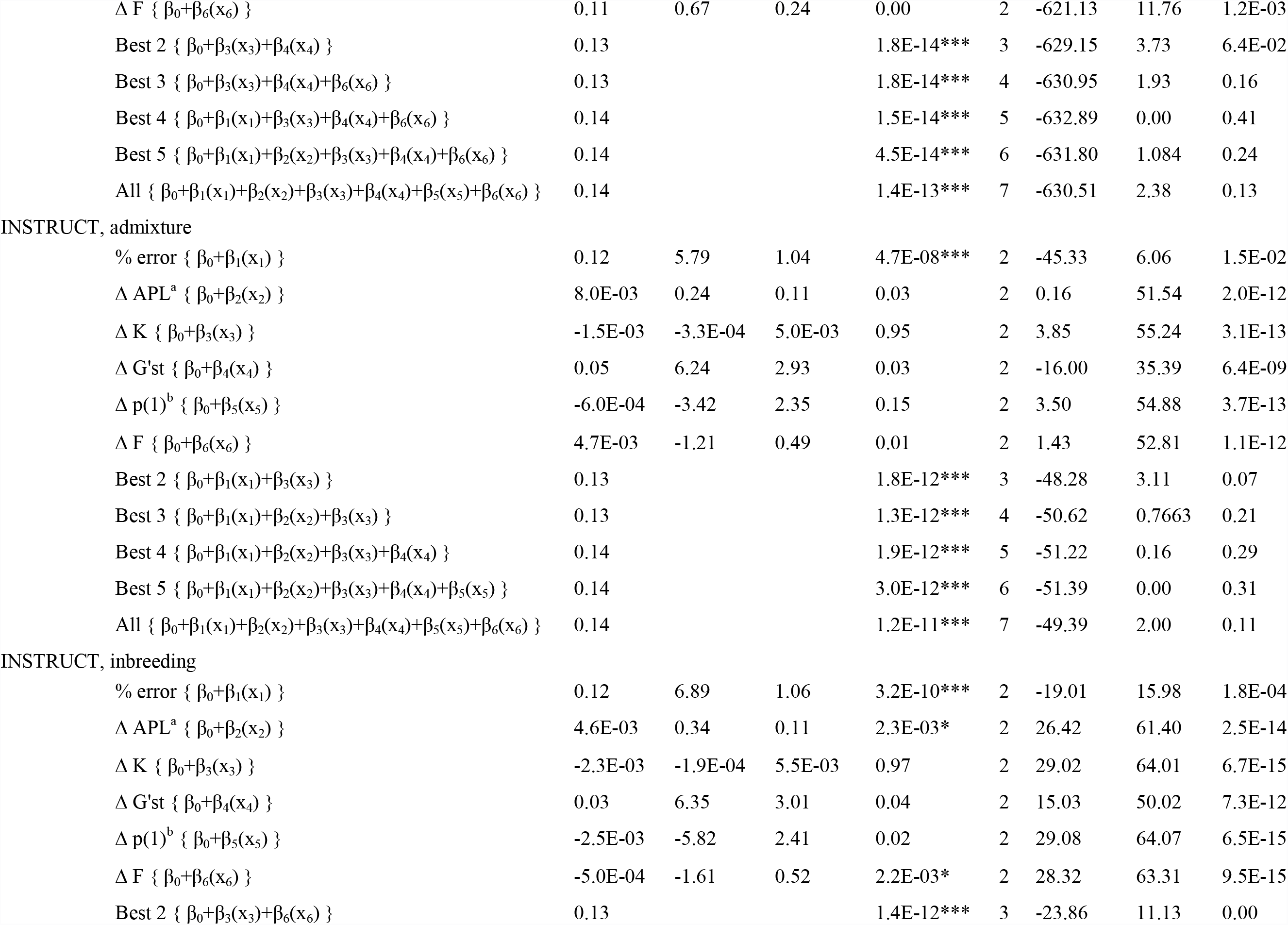
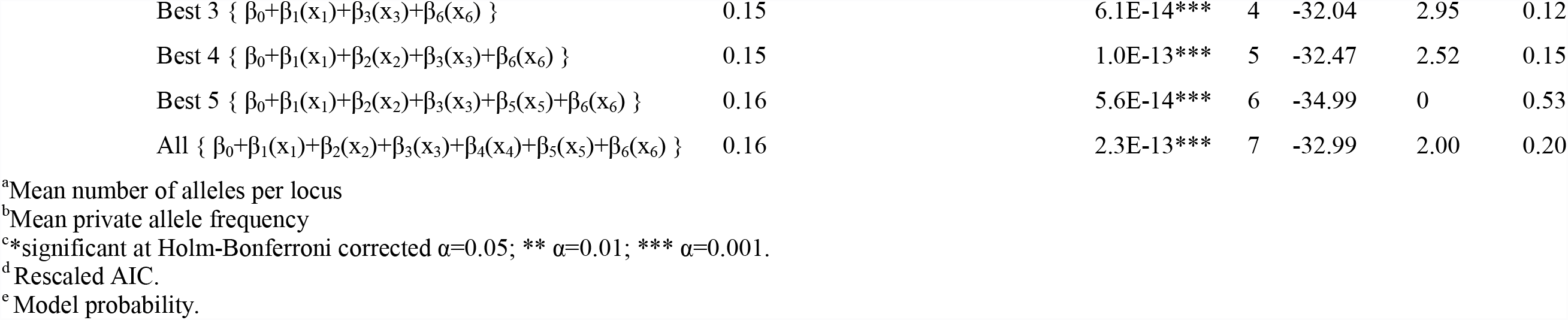
Multiple regression and likelihood analysis of six competing model effects explaining accuracy of model-based clustering methods subject to erroneous data, measured with the partition distance ratio metric.

| Method | Model | $R^2_{adj}$ | $\beta$ | SE | $p^c$ | K | AIC | $\Delta_i^d$ | $w_i^e$ |
| --- | --- | --- | --- | --- | --- | --- | --- | --- | --- |
| STRUCTURAMA |  |  |  |  |  |  |  |  |  |
| | % error { $\beta_0+\beta_1(x_1)$ } | 0.04 | -0.78 | 0.72 | 0.28 | 2 | -245.03 | 83.80 | 2.4E-19 |
| | $\Delta$ APL<sup>a</sup> { $\beta_0+\beta_2(x_2)$ } | 6.8E-03 | -0.01 | 0.05 | 0.84 | 2 | -238.05 | 90.78 | 7.3E-21 |
| | $\Delta$ K { $\beta_0+\beta_3(x_3)$ } | 0.43 | 0.12 | 0.01 | 1.2E-15*** | 2 | -318.23 | 10.60 | 1.9E-03 |
| | $\Delta$ G'st { $\beta_0+\beta_4(x_4)$ } | 0.05 | -1.94 | 3.15 | 0.54 | 2 | -246.64 | 82.19 | 5.4E-19 |
| | $\Delta$ p(1)<sup>b</sup> { $\beta_0+\beta_5(x_5)$ } | 0.01 | -0.13 | 1.37 | 0.92 | 2 | -240.54 | 88.29 | 2.6E-20 |
| | $\Delta$ F { $\beta_0+\beta_6(x_6)$ } | 0.15 | 1.51 | 0.56 | 7.6E-03 | 2 | -261.90 | 66.93 | 1.1E-15 |
| | Best 2 { $\beta_0+\beta_3(x_3)+\beta_6(x_6)$ } | 0.48 | | | 1.9E-20*** | 3 | -328.83 | 0 | 0.38 |
| | Best 3 { $\beta_0+\beta_1(x_1)+\beta_3(x_3)+\beta_6(x_6)$ } | 0.48 | | | 6.8E-20*** | 4 | -328.75 | 0.07 | 0.37 |
| | Best 4 { $\beta_0+\beta_1(x_1)+\beta_3(x_3)+\beta_4(x_4)+\beta_6(x_6)$ } | 0.48 | | | 4.0E-19*** | 5 | -327.20 | 1.63 | 0.17 |
| | Best 5 { $\beta_0+\beta_1(x_1)+\beta_2(x_2)+\beta_3(x_3)+\beta_4(x_4)+\beta_6(x_6)$ } | 0.47 | | | 2.4E-18*** | 6 | -325.24 | 3.59 | 0.06 |
| | All { $\beta_0+\beta_1(x_1)+\beta_2(x_2)+\beta_3(x_3)+\beta_4(x_4)+\beta_5(x_5)+\beta_6(x_6)$ } | 0.47 | | | 1.3E-17*** | 7 | -323.25 | 5.58 | 0.02 |
| INSTRUCT, no admixture |  |  |  |  |  |  |  |  |  |
| | % error { $\beta_0+\beta_1(x_1)$ } | 0.06 | 1.32 | 0.52 | 0.01 | 2 | -597.36 | 35.52 | 8.0E-09 |
| | $\Delta$ APL<sup>a</sup> { $\beta_0+\beta_2(x_2)$ } | -1.5E-03 | 0.10 | 0.05 | 0.07 | 2 | -567.72 | 65.17 | 2.9E-15 |
| | $\Delta$ K { $\beta_0+\beta_3(x_3)$ } | -9.0E-04 | -1.3E-03 | 1.4E-03 | 0.36 | 2 | -567.98 | 64.91 | 3.3E-15 |
| | $\Delta$ G'st { $\beta_0+\beta_4(x_4)$ } | 0.10 | 2.70 | 1.45 | 0.06 | 2 | -616.39 | 16.50 | 1.1E-04 |
| | $\Delta$ p(1)<sup>b</sup> { $\beta_0+\beta_5(x_5)$ } | 7.7E-03 | -1.01 | 1.21 | 0.40 | 2 | -571.96 | 60.93 | 2.4E-14 |

|  |  |  |  |  |  |  |  |  |
| --- | --- | --- | --- | --- | --- | --- | --- | --- |
| $\Delta F \{ \beta_0 + \beta_6(x_6) \}$ | 0.11 | 0.67 | 0.24 | 0.00 | 2 | -621.13 | 11.76 | 1.2E-03 |
| Best 2 $\{ \beta_0 + \beta_3(x_3) + \beta_4(x_4) \}$ | 0.13 | | | 1.8E-14*** | 3 | -629.15 | 3.73 | 6.4E-02 |
| Best 3 $\{ \beta_0 + \beta_3(x_3) + \beta_4(x_4) + \beta_6(x_6) \}$ | 0.13 | | | 1.8E-14*** | 4 | -630.95 | 1.93 | 0.16 |
| Best 4 $\{ \beta_0 + \beta_1(x_1) + \beta_3(x_3) + \beta_4(x_4) + \beta_6(x_6) \}$ | 0.14 | | | 1.5E-14*** | 5 | -632.89 | 0.00 | 0.41 |
| Best 5 $\{ \beta_0 + \beta_1(x_1) + \beta_2(x_2) + \beta_3(x_3) + \beta_4(x_4) + \beta_6(x_6) \}$ | 0.14 | | | 4.5E-14*** | 6 | -631.80 | 1.084 | 0.24 |
| All $\{ \beta_0 + \beta_1(x_1) + \beta_2(x_2) + \beta_3(x_3) + \beta_4(x_4) + \beta_5(x_5) + \beta_6(x_6) \}$ | 0.14 | | | 1.4E-13*** | 7 | -630.51 | 2.38 | 0.13 |
| INSTRUCT, admixture |  |  |  |  |  |  |  |  |
| % error $\{ \beta_0 + \beta_1(x_1) \}$ | 0.12 | 5.79 | 1.04 | 4.7E-08*** | 2 | -45.33 | 6.06 | 1.5E-02 |
| $\Delta APL^a \{ \beta_0 + \beta_2(x_2) \}$ | 8.0E-03 | 0.24 | 0.11 | 0.03 | 2 | 0.16 | 51.54 | 2.0E-12 |
| $\Delta K \{ \beta_0 + \beta_3(x_3) \}$ | -1.5E-03 | -3.3E-04 | 5.0E-03 | 0.95 | 2 | 3.85 | 55.24 | 3.1E-13 |
| $\Delta G'st \{ \beta_0 + \beta_4(x_4) \}$ | 0.05 | 6.24 | 2.93 | 0.03 | 2 | -16.00 | 35.39 | 6.4E-09 |
| $\Delta p(1)^b \{ \beta_0 + \beta_5(x_5) \}$ | -6.0E-04 | -3.42 | 2.35 | 0.15 | 2 | 3.50 | 54.88 | 3.7E-13 |
| $\Delta F \{ \beta_0 + \beta_6(x_6) \}$ | 4.7E-03 | -1.21 | 0.49 | 0.01 | 2 | 1.43 | 52.81 | 1.1E-12 |
| Best 2 $\{ \beta_0 + \beta_1(x_1) + \beta_3(x_3) \}$ | 0.13 | | | 1.8E-12*** | 3 | -48.28 | 3.11 | 0.07 |
| Best 3 $\{ \beta_0 + \beta_1(x_1) + \beta_2(x_2) + \beta_3(x_3) \}$ | 0.13 | | | 1.3E-12*** | 4 | -50.62 | 0.7663 | 0.21 |
| Best 4 $\{ \beta_0 + \beta_1(x_1) + \beta_2(x_2) + \beta_3(x_3) + \beta_4(x_4) \}$ | 0.14 | | | 1.9E-12*** | 5 | -51.22 | 0.16 | 0.29 |
| Best 5 $\{ \beta_0 + \beta_1(x_1) + \beta_2(x_2) + \beta_3(x_3) + \beta_4(x_4) + \beta_5(x_5) \}$ | 0.14 | | | 3.0E-12*** | 6 | -51.39 | 0.00 | 0.31 |
| All $\{ \beta_0 + \beta_1(x_1) + \beta_2(x_2) + \beta_3(x_3) + \beta_4(x_4) + \beta_5(x_5) + \beta_6(x_6) \}$ | 0.14 | | | 1.2E-11*** | 7 | -49.39 | 2.00 | 0.11 |
| INSTRUCT, inbreeding |  |  |  |  |  |  |  |  |
| % error $\{ \beta_0 + \beta_1(x_1) \}$ | 0.12 | 6.89 | 1.06 | 3.2E-10*** | 2 | -19.01 | 15.98 | 1.8E-04 |
| $\Delta APL^a \{ \beta_0 + \beta_2(x_2) \}$ | 4.6E-03 | 0.34 | 0.11 | 2.3E-03* | 2 | 26.42 | 61.40 | 2.5E-14 |
| $\Delta K \{ \beta_0 + \beta_3(x_3) \}$ | -2.3E-03 | -1.9E-04 | 5.5E-03 | 0.97 | 2 | 29.02 | 64.01 | 6.7E-15 |
| $\Delta G'st \{ \beta_0 + \beta_4(x_4) \}$ | 0.03 | 6.35 | 3.01 | 0.04 | 2 | 15.03 | 50.02 | 7.3E-12 |
| $\Delta p(1)^b \{ \beta_0 + \beta_5(x_5) \}$ | -2.5E-03 | -5.82 | 2.41 | 0.02 | 2 | 29.08 | 64.07 | 6.5E-15 |
| $\Delta F \{ \beta_0 + \beta_6(x_6) \}$ | -5.0E-04 | -1.61 | 0.52 | 2.2E-03* | 2 | 28.32 | 63.31 | 9.5E-15 |
| Best 2 $\{ \beta_0 + \beta_3(x_3) + \beta_6(x_6) \}$ | 0.13 | | | 1.4E-12*** | 3 | -23.86 | 11.13 | 0.00 |

|  |  |  |  |  |  |  |
| --- | --- | --- | --- | --- | --- | --- |
| Best 3 { $\beta_0 + \beta_1(x_1) + \beta_3(x_3) + \beta_6(x_6)$ } | 0.15 | 6.1E-14*** | 4 | -32.04 | 2.95 | 0.12 |
| Best 4 { $\beta_0 + \beta_1(x_1) + \beta_2(x_2) + \beta_3(x_3) + \beta_6(x_6)$ } | 0.15 | 1.0E-13*** | 5 | -32.47 | 2.52 | 0.15 |
| Best 5 { $\beta_0 + \beta_1(x_1) + \beta_2(x_2) + \beta_3(x_3) + \beta_5(x_5) + \beta_6(x_6)$ } | 0.16 | 5.6E-14*** | 6 | -34.99 | 0 | 0.53 |
| All { $\beta_0 + \beta_1(x_1) + \beta_2(x_2) + \beta_3(x_3) + \beta_4(x_4) + \beta_5(x_5) + \beta_6(x_6)$ } | 0.16 | 2.3E-13*** | 7 | -32.99 | 2.00 | 0.20 |

<sup>a</sup>Mean number of alleles per locus
<sup>b</sup>Mean private allele frequency
<sup>c</sup>\*significant at Holm-Bonferroni corrected $\alpha=0.05$ ; \*\* $\alpha=0.01$ ; \*\*\* $\alpha=0.001$ .
<sup>d</sup>Rescaled AIC.
<sup>e</sup>Model probability.

